# Effect of fire and thinning on fine-scale genetic structure and gene flow in fire-suppressed populations of sugar pine (*Pinus lambertiana* Douglas)

**DOI:** 10.1101/448522

**Authors:** Brandon M. Lind, Malcolm P. North, Patricia E. Maloney, Andrew J. Eckert

## Abstract

Historically, frequent, low-severity fires in dry western North American forests were a major driver of ecological patterns and processes, creating resilient ecosystems dominated by widely-spaced pine species. However, a century of fire-suppression has caused overcrowding, altering forest composition to shade-tolerant species, while increasing competition and leaving trees stressed and susceptible to pathogens, insects, and high-severity fire. Exacerbating the issue, fire incidence is expected to increase with changing climate, while fire season has been observed to begin earlier and last longer than historic trends. Forest thinning and prescribed fire have been identified as important management tools to mitigate these risks. Yet little is known of how thinning, fire, or their interaction affect contemporary evolutionary processes of constituent pine species that influence fitness and play an important role in the opportunity for selection and population persistence. We assessed the impact of widely used fuel reduction treatments and prescribed fire on fine-scale gene flow on an ecologically important and historically dominant shade-intolerant pine species of the Sierra Nevada, *Pinus lambertiana* Dougl. Treatment prescription (no-thin-no-fire, thin-no-fire, and fire-and-thin) was found to differentially affect both fine-scale spatial and genetic structure as well as effective gene flow in this species. Specifically, the thin-no-fire prescription increases genetic structure (spatial autocorrelation of relatives) between adults and seedlings, while seed and pollen dispersal increase and decrease, respectively, as a function of increasing disturbance intensity. While these results may be specific to the stands at our study site, they indicate how assumptions relating to genetic effects based on spatial structure can be misleading. It is likely that these disequilibrated systems will continue to evolve on unknown evolutionary trajectories. The long-term impacts of management practices on reduced fitness from inbreeding depression should be continually monitored to ensure resilience to increasingly frequent and severe fire, drought, and pest stresses.

## Introduction

Many aspects of conifer biology are affected by a tree’s surrounding environment as well as the density of hetero- and conspecifics. For instance, outcrossing rates of conifer species are often tied to population density (Farris & Mitton 1984) and surrounding tree heights (O’Connell *et al.* 2003), while removal of proximal individuals can increase pollen and gene flow distances by reducing potential mates and removing once impeding vegetation. Thus, disturbance, *sensu lato*, has the potential to alter contemporary demographic and reproductive dynamics through both direct (population-level) and indirect (ecological-level) impacts (Mouillot *et al.* 2013).

Historically, natural disturbances such as fire were commonplace and equilibrated many ecosystem functions and processes in forests of the western United States (Covington *et al.* 1994). Fire regimes in these regions had return intervals on decadal scales (10-17 years; North et al. 2005), in contrast to wetter climates where fire return intervals were (sub)centennial (50+ years [North *et al.* 2016]). Resultantly, these ecosystems experienced frequent, low-severity burns and were populated by fire-adapted species, creating forests dominated by resilient, widely spaced pine trees. Yet over the past 150 years, anthropogenic influence has resulted in forests that are now fire-suppressed and overgrown by shade-tolerant species, causing increased competition, leaving trees stressed and susceptible to fungal and bark beetle attacks (Bonello *et al.* 2006).

Stand densification has also increased the frequency and probability of contemporary, high-severity fires. Between 2012 and 2014 in California alone, 14,340 fires burned 1.1 million acres and injured or killed nearly 300 individuals (NIFC 2014). Collectively, fires across California, the Great Basin, Southwest, and Rocky Mountain territories have burned a combined 8.8 million acres between 2014 and 2015 (NIFC 2015), while Forest Service scientists predict future fires to reach unprecedented levels, covering over 12-15 million acres annually (USDA Forest Service 2016a) requiring the United States Forest Service (USFS) to budget $2,300,000,000 on wildfire management, suppression, and preparedness for the 2016 fiscal year (USDA Forest Service 2016b). Exacerbating the issue, analyses of fire season length and onset have shown that seasons are beginning earlier and lasting longer than historic trends (Westerling 2006) while climate models predict extreme weather favorable to fire to become more frequent, and ignited fires to increase in severity, size, and required suppression efforts (Miller *et al.* 2009).

Because of these contemporaneous trends, large-scale forest thinning projects have been implemented to simultaneously restore fire-frequent ecosystems to their pre-settlement resilience as well as to protect urban development and human life, as fuel reduction treatments have been shown to be an effective tool in decreasing fire severity and ignition probability (Agee & Skinner 2005; Schwilk *et al.* 2009; Safford *et al.* 2009). These thinning treatments are often applied by determining DBH thresholds for cutting, and in some cases the density of leave-trees as well to reduce overall fuel load and continuity. For example, the Sierra Nevada Forest Plan Amendment (USDA Forest Service 2004) mandates that 50% of initial understory thinning treatments take place near urban populations, while the remaining thinning take place in natural wildland stands. To encourage fire resiliency the USFS has implemented fuel reduction treatments across 6.1 million acres of western, fire-suppressed forestland in 2014 (USDA Forest Service 2016a). Further, forest and fire scientists are calling for an overhaul of management policy to implement these thinning treatments to a far greater extent (North *et al.* 2015). While congruent with historic forest structure, these actions will orient these already disequilibrated systems on trajectories of unknown evolutionary consequence.

Through timber harvests, land use conversion, and fire suppression, forests have undergone systemic shifts in composition, structure, and disturbance regimes that are incongruous to the natural and evolutionary histories of endemic species (Collins *et al.* 2011; Larson & Churchill 2012). Consequentially, anthropogenic forest disturbance has been at the forefront of conservation attention for decades (Ledig 1988; 1992). The extent of human impact on forested land has received particular attention as a result of the empirical expectations developed from population genetic theory. Specifically, because of the reduction in individual tree density overall, and in particular for larger trees that asymmetrically contribute gametes to reproduction (Richardson *et al.* 2014), harvested forests are thought to be specifically subjected to population bottlenecks, potentially altering existing mating systems or available gene pools while decreasing genetic variability within populations and increasing differentiation from native stands (Smouse *et al*. 2001, Cloutier *et al.* 2006; Kramer *et al.* 2008; Lowe *et al.* 2015). These consequences can influence the fitness of affected populations, as drastic changes in gene pool availability or mating system alter a population’s potential to adapt to local conditions, and inbreeding depression can have deleterious effects on growth and reproductive output (e.g., reproductive capacity or rates of embryo abortion; Williams & Savolainen 1996; Sorensen 2001; Savolainen et al. 2007; Savolainen & Pyhäjärvi 2007).

Past studies investigating the genetic effects of North American forest management show mixed evidence of harvest influence. These studies often sub-sample populations and primarily focus on diversity consequences across a range of molecular markers (often microsatellites). Many management studies of North American conifers compare genotypic diversity indices (e.g., *H*_*E*_, *H*_*O*_, allelic richness, etc.) between treatments to detect management influence (Cheliak *et al.* 1988; Gömöry 1992; Buchert *et al.* 1997; Adams *et al.* 1998; Rajora *et al.* 2000; Macdonald *et al.* 2001; Perry & Bousquet 2001; Rajora & Pluhar 2003; El-Kassaby *et al.* 2003; Marquardt *et al.* 2007; Fageria & Rajora 2013a; b). However, the same diversity values can manifest under completely different scenarios and tests of significance between population values for a small number of markers may therefore be under-informative, particularly for sub-sampled populations, as these differences can result from sampling bias or from evolutionary processes unrelated to management. Additionally, these investigations also often employ *F*_*ST*_ analyses to assess statistical significance between treated and untreated stands (Thomas *et al.* 1999; Perry & Bousquet 2001; Marquardt *et al.* 2007; Fageria & Rajora 2013a; b). Though when used in this context, this test is simply signifying whether the allelic frequencies in (sub)populations under study are likely to have been sampled from the same ancestral population (Holsinger & Weir 2009). Very often, the treated and untreated stands are physically adjacent (derived of a common ancestral population) and only under extreme perturbation should significance be expected. In cases where significance is detected, and other than to assess relative diversity between stands, such differentiation does little to inform how management is affecting ongoing evolutionary processes affecting fitness, as such processes may ameliorate bottlenecks due to management. It would therefore be difficult to draw such conclusions without assessing other stand and evolutionary dynamics.

Very seldom in North American studies of forest management are evolutionary processes influencing fitness specifically examined (but see Neale & Adams 1985). Yet when studies are done and nonsignificant findings are found, authors generally caution interpretation (Finkeldey & Ziehe 2004; Namroud *et al.* 2012). Very often the scale of sampling (both in terms of numbers and spatial extent of individuals and the degree of temporal variation), as well as the lack of investigation into evolutionary dynamics have been offered as inadequate, and that further investigation into evolutionary consequences of natural and anthropogenic disturbance could give valuable insight to forest managers and fill a vital knowledge gap in this regard (Namroud *et al.* 2012). Indeed, incongruence between theoretical predictions and empirical results from studies evaluating genetic consequences of forest disturbance has created a paradox within the literature (Kramer *et al.* 2008). Yet as Lowe et al. (2015) point out, we may have been looking in the wrong place. They argue that instead of simply assaying mature cohorts to understand the genetic consequences of disturbance, future attention should include progeny arrays as well as the relative regenerative success across a wide range of influences. Additionally, they contend that the type and magnitude of the genetic response itself may be better understood through the variation in mating and breeding systems of studied species. Of particular importance, Lowe *et al.* (2015) advise scientists that the most fruitful research endeavors will incorporate quantitative approaches to understanding evolutionary mechanisms, specifically those connecting changes in pollination to mating systems and evolutionary fitness, and that these efforts will likely generate critical knowledge regarding the mechanisms driving the dynamics we observe.

Interactions between fire and forest thinning management are certain. To ensure forests are resilient to frequent fire and disturbance, and provide habitat for public recreation and native wildlife, the interactive impact of management and fire must be understood in an evolutionary framework. Here, we investigated the evolutionary impact of forest management on fire-suppressed populations of the historically dominant and ecologically important sugar pine (*Pinus lambertiana* Dougl.) within Teakettle Experimental Forest (TEF), a USFS site located in the central Sierra Nevada of California. Using microsatellite markers, we employ parentage analysis and assess impact upon various processes known to affect fitness such as mating patterns, effective dispersal distances, and fine-scale (<300m) genetic structure. We tested hypotheses relating to effective dispersal distances of pollen and seed, the relative spatial and genetic structure of tree classes (size/age and species) within treatments, and for differences in these measures across treatments. Although the genetic structure of adults is due to an interaction between the evolutionary history of the stand and the applied treatment, mating patterns and seedling recruitment will determine long-term impacts of management. Our results show that thinning alone increases fine-scale genetic structure (i.e., spatial autocorrelation of relatives), and that the majority of pollen and seed dispersal take place at the same scale. While effects of such treatments will vary by location, the degree of thinning and the choice of leave-trees should be tailored to a given stand, and spatial structure (arrangement of individuals across the landscape) should not be conflated with spatial genetic structure (arrangement of relatives across the landscape). By avoiding treatments that increase genetic structure, managers may be able to decrease seed abortion due to inbreeding and thus increase effective seed rain of species with management importance.

## Methods

### Study area, sampling, and focal species

Teakettle Experimental Forest (TEF) is a fire-suppressed, old-growth forest watershed in the central Sierra Nevada mountains of California. The 1300-ha watershed ranges from 1900– 2600m in elevation and consists of five conifer species: white fir (*Abies concolor* [Gordon] Lindley ex Hildebrand), red fir (*A. magnifica* A. Murray), incense cedar (*Calocedrus decurrens* [Torr] Florin), Jeffrey pine (*Pinus jeffreyi* Balf.), and sugar pine (*P. lambertiana*). Historically, fire burned the area every 11-17 years, but has been suppressed for 135 years (North *et al.* 2005) while logging had been completely absent (North et al. 2002). Six treatments were applied to neighboring 4-ha plots (each 200m x 200m, Figure 1a) by crossing two levels of burn (no-fire and fire) with three levels of thinning (no-thinning, overstory-thinning, and understory-thinning). The understory thinning prescription followed guidelines in the California spotted owl (CASPO) report (Verner *et al.* 1992), which is now widely used for fuel management in California (SNFPA 2004). We therefore focus our analyses on untreated stands and those treated with or in combination with this understory thinning treatments (see below). Each treatment was replicated three times (18 plots covering 72ha). Understory-thinning removed all trees with a diameter at breast height (DBH) ≤76cm and ≥25cm, while overstory-thinning removes all trees >25cm DBH except 18-22 of the largest trees per hectare. Treatments were applied over 2000 and 2001. Plot inventories of pre-treatment (1999), and post-treatment (2004 and 2011) conditions mapped individual trees on a 3D coordinate system (colored dots, Figure 1b). Only standing boles ≥5cm DBH were included in plot inventories, which recorded species, DBH, spatial coordinates, decay class, and forest health metrics (e.g., presence/absence of insects and pathogens). Post-treatment inventories updated these metrics, and added individuals to the dataset once they reached 5cm DBH. Here, seedling and saplings are all pine stems <5cm DBH. For these, basal diameter and spatial coordinates were recorded over the summers of 2012 and 2013 while collecting needle tissue samples from the full census of all live *P. lambertiana* (*N* = 3,135). *Pinus lambertiana* is a historically dominant member of mixed-conifer forests of the Sierra Nevada, and continues to play important ecological roles. This species is shade-intolerant and is an important focus of restoration in the Sierra Nevada range.

**Figure 1.**
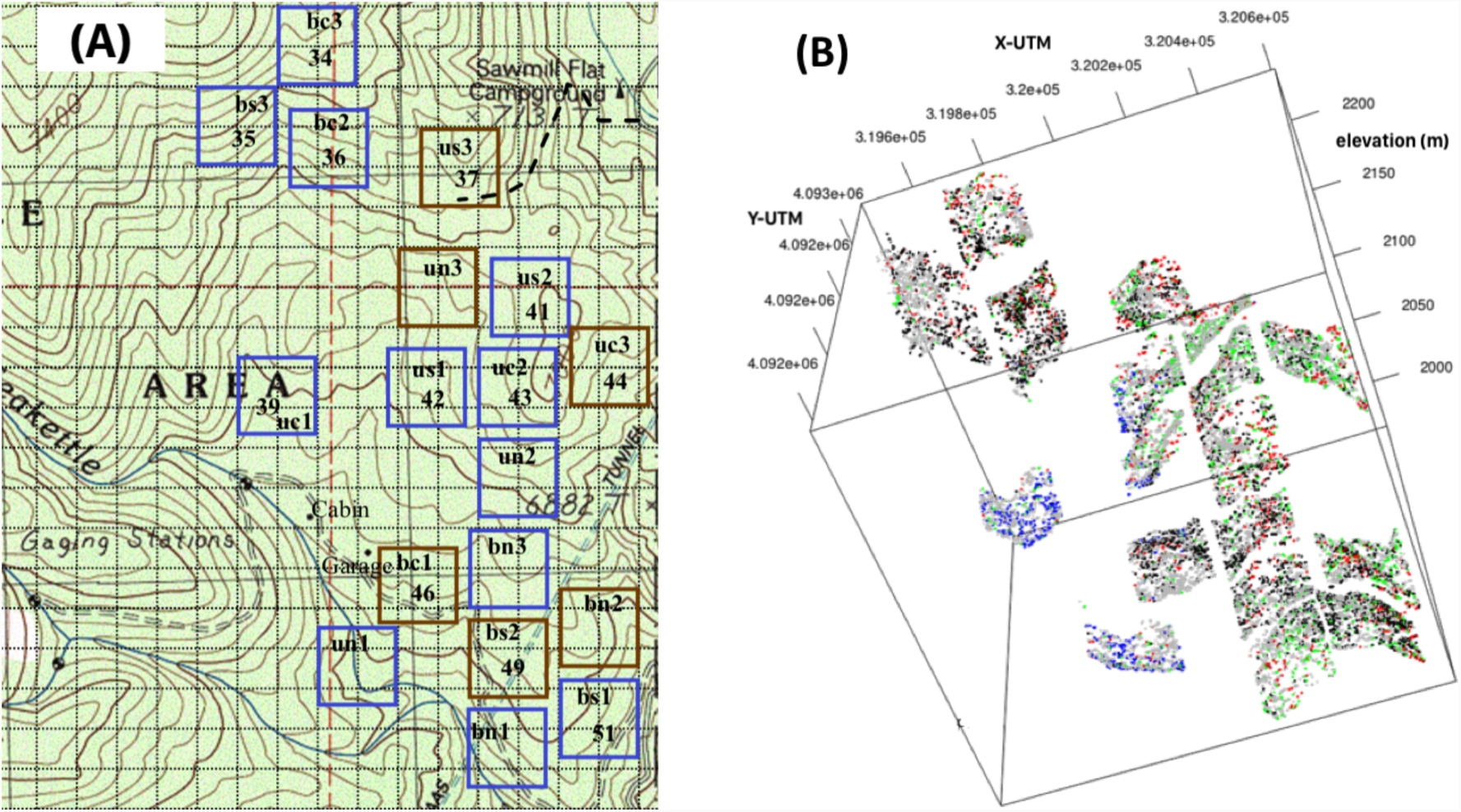
Teakettle Experimental Forest, California (Latitude: 36.9606, Longitude: -119.0258). (A) Topographic map and spatial arrangement of treatments (BC = burned understory thin; BN = burned no-thin; BS = burned shelterwood thin; UC = unburned understory thin; UN = unburned no-thin; US = unburned shelterwood thin). Replicates for each treatment are numbered one through three from south to north. (B) Mapped coordinates (Universal Transverse Mercator) and elevation (meters) of pre-treatment adults ≥ 5cm diameter at breast height. (green: P. lambertiana, red: P. jeffreyi, gray: A. concolor, blue: A. magnifica, orange: C. decurrens, black: Quercus, Salix, and remaining species.).

### Analysis of tree spatial structure

Using plot-level *P. lambertiana* individuals, we estimated spatial structure of seedlings and adults across 10-meter distance classes, *r*, separately using univariate inhomogeneous pair correlation functions (*g*_inhom_(*r*)) from the `spatstat` library (Baddeley *et al.* 2015) with an isotropic edge correction. This statistic was chosen over Ripley’s K, or its linearized version (*L*), because of advocacy for *g*_inhom_(*r*) over these statistics (see `spatstat` manual). This analysis tests the null hypothesis that the 2D spatial arrangement of points (adults or seedlings) is not significantly different from complete spatial randomness (CSR; i.e., a Poisson distribution of inter-point distances with *g*_inhom_ ogeneous intensities of points), where support for the alternative hypothesis is indicative of ecological factors driving spatial patterning. We calculated null confidence envelopes for each test using 199 null simulations of CSR using the same intensity of the pattern of individuals analyzed (equivalent to an alpha value of 0.01; see `spatstat` manual). For trees that coincide with the null model of CSR, *g*_inhom_(*r*) = 1, with spatial aggregation *g*_inhom_(*r*) > 1, and with spatial inhibition *g*_inhom_(*r*) < 1 (Baddeley *et al*. 2015); significance was judged using the null confidence envelopes. We repeated this analysis for the dominant shade-tolerant individuals (all *Abies* individuals). We extended the univariate function to its bivariate equivalent, *g*_inhom_(*r*), to test for spatial affinity between two groups *i* and *j*, using similar methods as above for edge correction and null confidence envelopes. We calculated *g*_*inhom,i,j*_(*r*) between unique combinations of *P. lambertiana* adults, *P. lambertiana* seedlings, and shade-tolerant *Abies* individuals. Hypothesis testing and interpretation of bivariate *g*_*inhom,i,j*_(*r*) was carried out as with univariate *g*_inhom_(*r*). Results from these analyses allow comparison of standing spatial structure against spatial genetic autocorrelation (see below), to make inferences about the ecology of these species, and how treatments at TEF are affecting ongoing evolutionary dynamics.

### DNA extraction, microsatellite amplification

Total genomic DNA was extracted according to manufacturer’s protocol using the DNeasy 96 Plant Kit (Qiagen, Germantown, MD) from *P. lambertiana* samples within a subset of the factorial treatments at TEF: unburned-no-thin control plots (hereafter UN), understory-thin (CASPO) plots without burn application (hereafter UC), and burned understory-thin plots (hereafter BC) for a total of 1,348 individuals. Herein, we often refer to patterns across these treatments in terms of increasing disturbance intensity (i.e., from UN to UC to BC). Three chloroplast (paternally inherited, Wofford *et al.* 2014: pt71936, pt87268, pc10) and four nuclear (biparental inheritance, Echt *et al.* 1996: rps50, rps02, rps12, rps39) microsatellite markers were amplified (using fluorescent dyes NED, PET, VIC, and FAM) per the original publications with minor modifications using BIO-RAD iProof high fidelity DNA polymerase (see Supplemental Information). The chloroplast markers were chosen for their primer conservation across *Pinus*, *Trifoliae*, *Parrya*, and *Quinquifolia* subsections of the *Pinus* genus (Wofford *et al.* 2014) while the chosen nuclear markers have been amplified in eastern white pine (*P. strobus* L., Echt *et al.* 1996) and both sets successfully amplified on a subset of individuals at TEF judged by gel electrophoresis. Multiplexed individuals (one fluorescent dye per well) were analyzed using the Applied Biosystems 3730xl fragment analyzer at Cornell University (http://www.biotech.cornell.edu/brc/genomics-facility) and genotypes were called using GeneMaker v2.6.7 (see Supplemental Info; http://www.softgenetics.com/GeneMarker.php).

### Genetic diversity measures

Treatment-specific diversity measures (total number of alleles, *A*_T_; mean number of alleles per locus, *A*; effective number of alleles per locus, *A*_e_; observed and expected heterozygosity for nuclear markers, respectfully *H*_*o*_, *H*_e_; average number of private alleles, *A*_P_; and overall means for each category) were calculated for each treatment and averaged across loci in order to compare dynamics at TEF to published studies. For estimates of *H*_*o*_ and *H*_e_, only nuclear markers were used. To quantify variation we report standard deviation. We calculated hierarchical multi-locus *F*_ST_ (Weir & Cockerham 1984) for nuclear markers using the `hierfstat` package (Goudet & Jombart 2015) and treatment-specific *F*_ST_ to compare across treatments. Single- and multi-locus exclusion probabilities for parentage analysis (see below) were calculated using python scripts modified from `gstudio` (v1.5.0; Dyer 2016).

### Analysis of spatial genetic structure

To quantify spatial genetic autocorrelation at a distance class *h* (hereafter 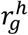), we used multi-locus genetic distances (Smouse & Peakall 1999) and Euclidean geographic distances among spatial coordinates of individuals across distances classes *h* corresponding to approximately 10-meter bins for *P. lambertiana* seedlings, *P. lambertiana* adults, as well as a bivariate approximation for the clustering of *P. lambertiana* adult genotypes to those of seedlings. For a distance class, *h*, spatial patterning of multi-locus genotypes are unrelated to (i.e., random relative to) the spatial patterns of individuals if 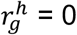, aggregated if 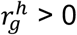, and dispersed if 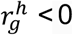. We estimated null confidence intervals by taking the 2.5^th^ and 97.5^th^ quantiles of *M* = 1000 estimates of 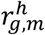, where 999 of these estimates were computed by randomly permuting individual genotypes across empirical spatial coordinates, with the *M*^th^ permutation being the empirical estimate of 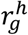 itself (Smouse & Peakall 1999). Using the `PopGenReport` package (Adamack & Gruber 2014) we created correlograms for nuclear and chloroplast markers both in isolation and in combination, but present only those using full genotypes as correlograms by marker type showed similar patterns as full genotypes. We used these correlograms to compare and contextualize treatment-level genetic structure with ongoing evolutionary dynamics such as that of fine-scale gene flow.

### Parentage analysis

To quantify fine-scale gene flow at TEF, we conducted parentage analysis using our genetic markers and spatial coordinates of individuals. Joint estimation of parentage and dispersal parameters of seed and pollen were achieved by expanding methods of Moran & Clark (2011). This method simultaneously estimates parentage and dispersal kernel parameters for seed and pollen within a Bayesian framework, taking into account genotyping error and variation in individual fecundity while treating dispersal processes inside and outside of the mapped areas in a coherent manner, which is critical if the dispersal kernel is to reflect both long- and short-distance movement. Here, all sampled adults are characterized by their genotype and mapped coordinate. Additionally, for seedlings there is also an estimated pedigree, which can consider any adult as either mother or father (though we excluded possible selfing events). The probability of the pedigree considering two sampled parents, before incorporating information regarding genotype, is estimated from the probability of pollen-to-mother movement over the given distance and of seed movement over the distance between mother and seedling, as well as the parental prior distribution for fecundity. Pollen production was considered proportional to fecundity (as in Moran & Clark 2011) and was estimated by fitting a 2^nd^-order power polynomial regression to data from Figure 6 in Fowells & Schubert (1956) where Cone Count = 0.0098(dbh^2^) - 0.4811(dbh) + 10.651. We then set fecundity for all adults <25cm DBH to zero given observed cone counts from Fowells & Schubert (1956). For dispersal priors, we set the seed dispersal kernel shape parameter, *u*_*s*_, to 253.31, (a mean dispersal distance of 25m; Millar *et al*. 1992; Fowells & Schubert 1956) and the pollen dispersal kernel shape parameter prior, *u*_p_, to 2279.72, (a mean pollen distance of 75m; Wright 1976; Neale 1983; Millar *et al*. 1992). For priors to the standard deviation of mean dispersal we set seed (pollen) to 1013.21 (9118.90) corresponding to standard deviations of 50m (75m).

Given that either parent could have produced the offspring, the likelihood that this pair is the true parents relative to all other possible parent pairs depends on the dispersal kernel priors for seed and pollen, and the seed and pollen production of all trees both inside and outside of the plot (the fraction of all possibilities; Moran & Clark 2011). To evaluate the probability of an offspring having one parent in the plot and the other outside of the plot, a set of potential out-of-plot parent-densities, *dp*_1_,…, *dp*_20_, each 10m progressively outside of the plot is considered (see figure S3.1 in Moran & Clark 2011). Pollen and seed movement into the plot is approximated by assuming first that all seed/pollen produced within each quarter-polygon, *v*, originates from a tree located *dp*_*v*_ meters from the midpoint of each side outside of the plot. The expected out-of-plot pollen (seeds) reaching an in-plot mother (a seedling’s location) from each quarter-polygon outside of the plot is calculated based on the average density and average fecundities of trees outside of the plot and then multiplied by the probability of dispersal to the point within the plot. Summing over each distance class gives the total expected out-of-plot pollen/seed dispersal to points inside of the plot. However, to calculate the probability of an in-plot versus an out-of-plot father, the expected pollen arriving at an out-of-plot mother from another out-of-plot father must first be calculated using the concentric polygons around the sampled plot and the distance classes described above. The fraction of rings falling outside the plot determines the fraction of pollen received from each distance class, *dp*_*v*_ expected to come from outside trees. Once error rates (*e*_1_) and dropout rates (*e*_2_) of genotyping are calculated through regenotyping individuals (see Supplemental Information), the probability of a pedigree, seed and dispersal parameters given the offspring genotype, distances, error rates, and pollen/seed production can be estimated (Moran & Clark 2011). Very rarely have previous studies investigating effects of forest management (or using parentage analysis towards such goals) incorporated error and dropout rates into subsequent inferences.

For the current study, out-of-plot densities were extrapolated from densities and DBH distributions (our proxy for fecundity) revealed in pre-treatment surveys (North et al. 2002). Due to the proximity of the treated plots, all adult trees were considered simultaneously for parentage assignment. Additionally, instead of considering any given pedigree as symmetrical (i.e., with no consideration for which tree was the pollen or seed donor) we utilize genotyped markers separately to consider whether a given pedigree is for a mother-father pair, or for a father-mother pair (i.e., we only considered nuclear markers for a potential mother, and all markers for a potential father). The most probable pedigree for each seedling was identified by assessing the proportion of the proposed pedigree across chains in the Gibbs sampler (as in Moran & Clark 2011), in which we used 500,000 steps and a burn-in of 30,000. This method was further modified to improve computational efficiency by multiprocessing appropriate elements of the script by utilizing custom python scripts and the `SNOW` library (v0.4-2; Tierney *et al*. 2016) in R (v3.3.3; R Core Team 2017). We replicated each run three times, and judged convergence within and between runs in R.

### Using parentage analysis to further quantify fine-scale gene flow

In addition to estimates of the mean seed and pollen dispersal (see above), we used these parentage assignments to further classify fine-scale gene flow at TEF. Using the full set of most probable pedigrees, we quantified the number of in-plot vs. out-of-plot dispersal events averaged across replicates for a given treatment. Then, using the most probable parentage assignment for each offspring, we quantified mean dispersal distances from sampled mothers to seedlings, and between sampled fathers to sampled mothers. To better account for uncertainty in parentage assignment (i.e., to account for fractional parentage assignment), we calculated mean dispersal distance by treatment by considering all pedigrees with known individuals weighted by the probability of assignment. Specifically, for mean seed dispersal, for each seedling we calculated the weighted average of mother-offspring distances across pedigrees of non-zero probability that included known mothers in the dataset. Each weight was the probability of assignment, *p*_*seed,pedigree*_, divided by the probability of assignment of this seedling to a known mother (1 - *U*_*M*_) where *U*_*M*_ is the sum of the probabilities across all non-zero pedigrees that included an unsampled mother. Treatment-level averages were then calculated across these weighted distances. For pollen dispersal, for each seedling we considered only pedigrees of non-zero probability where both the mother and father were known, weighting each distance by the probability of assignment, *p*_*seed,pedigree*_, divided by the probability of assignment to known parents (1-*U*_*seed,pedigree*_) where *U*_*seed,pedigree*_ is the sum of the probabilities across all non-zero pedigrees that included at least one unsampled parent. Treatment-level averages were then calculated from these weighted distances and significance was determined using a Kruskal-Wallis test with an alpha value of 0.05.

Scripts used in analyses described above can be found in IPython notebook format (Pérez & Granger 2007) at https://github.com/brandonlind/teakettle.

## Results

### Analysis of tree spatial structure

#### Univariate Analysis

Across treatments, *P. lambertiana* adults exhibited spatial aggregation at distance classes less than 20m, which decreased with increasing disturbance intensity with UN plots showing the greatest magnitudes of *g*_inhom_(*r*) at these small distance classes (Figure 2 first row). For adult shade-tolerant species (all *Abies* individuals), the extent of spatial aggregation at large distance classes decayed with increasing disturbance intensity (Figure 2 second row) where UC generally exhibited greater magnitudes of *g*_inhom_(*r*) than BC in small distance classes. For *P. lambertiana* seedlings, spatial structure was similar across treatments, though UN generally had significant aggregation and much larger magnitudes of *g*_inhom_(*r*) at larger distance classes than other treatments, while BC seedlings exhibited greater magnitudes of *g*_inhom_(*r*) across small distance classes than either UC or UN (Figure 2 third row).

**Figure 2.**
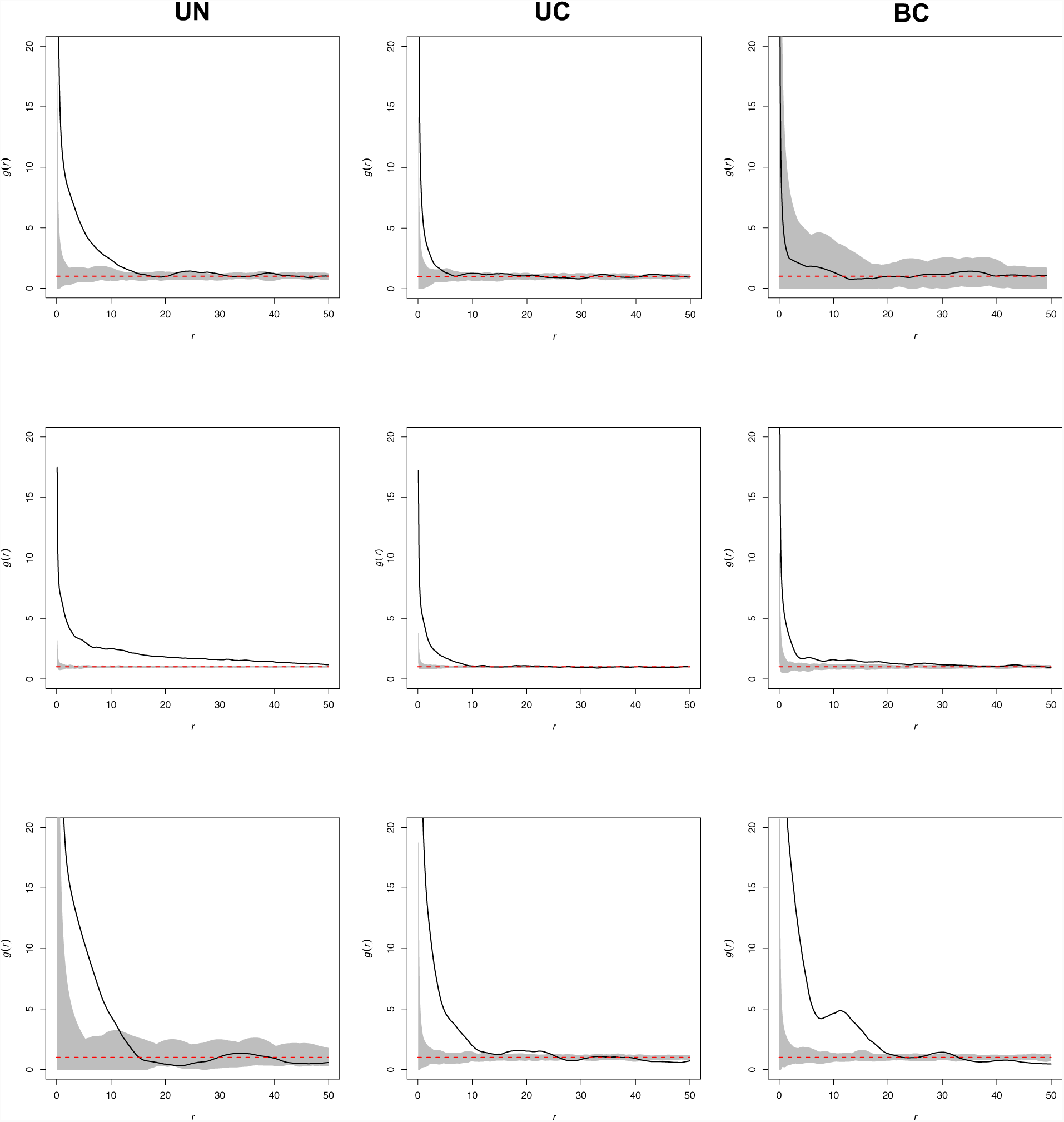
Representative figures of univariate analysis of spatial structure, *g*_inhom (*r*)_, by treatment replicate for each distance class, r. First row: adult P. lambertiana (PiLa); second row: adult shade tolerant (A. magnifica and A. concolor = ShadeTol); third row: P. lambertiana (PiLa) seedlings. Disturbance intensity increases by column from left to right. These figures show that with increasing disturbance intensity there is a diminution of the degree of spatial structure within classes. Gray : null confidence envelope; Solid black line : observed *g*_inhom_ (*r*). Red dashed line : null expectation of complete spatial randomness, *g*_inhom_ (*r*) = Individuals are aggregated if *g*_inhom_ (*r*) > 1, inhibited if *g*_inhom_ (*r*) < 1. See Supplemental Figures S1-S3 for all plots.

#### Bivariate Analysis

The spatial affinity of *P. lambertiana* seedlings to *P. lambertiana* adults, *g*_*inhom,seedling,adult*_(*r*), decreased with intensity of disturbance (i.e., from UN, to UC, to BC). UN plots showed consistent inhibition across distance classes greater than 15m, whereas UC plots tended to align with the lower extent of the confidence interval with fewer instances of significant inhibition (Figure 3). A similar trend for increasing spatial inhibition between *P. lambertiana* seedlings and shade-tolerant adults (*g*_*inhom,seedling,adult*_(*r*)), as well as for *P. lambertiana* adults and shade-tolerant adults (*g*_*inhom,PiLa–adult,shadetol*_(*r*); Figure 3) where UN generally had a greater inhibition than UC or BC, though BC exhibited evidence of spatial inhibition between groups. The results from the uni- and bivariate analyses of spatial patterns suggest that pines are generally clustered with other pines, shade-tolerant individuals are clustered with other shade-tolerant individuals, but shade-tolerant adults generally show spatial inhibition with pine individuals of both classes. Additionally, *P. lambertiana* seedlings showed similar clustering across all treatments, suggesting a similar pattern of response to the environment. Further, together with the univariate spatial clustering of *P. lambertiana* seedlings at small distance classes, these bivariate results suggest there may be ecological drivers influencing realized patterns of seedlings across microenvironments (e.g., perhaps sites with decreased competition for [or optimal levels of] water, nutrients, or light).

**Figure 3.**
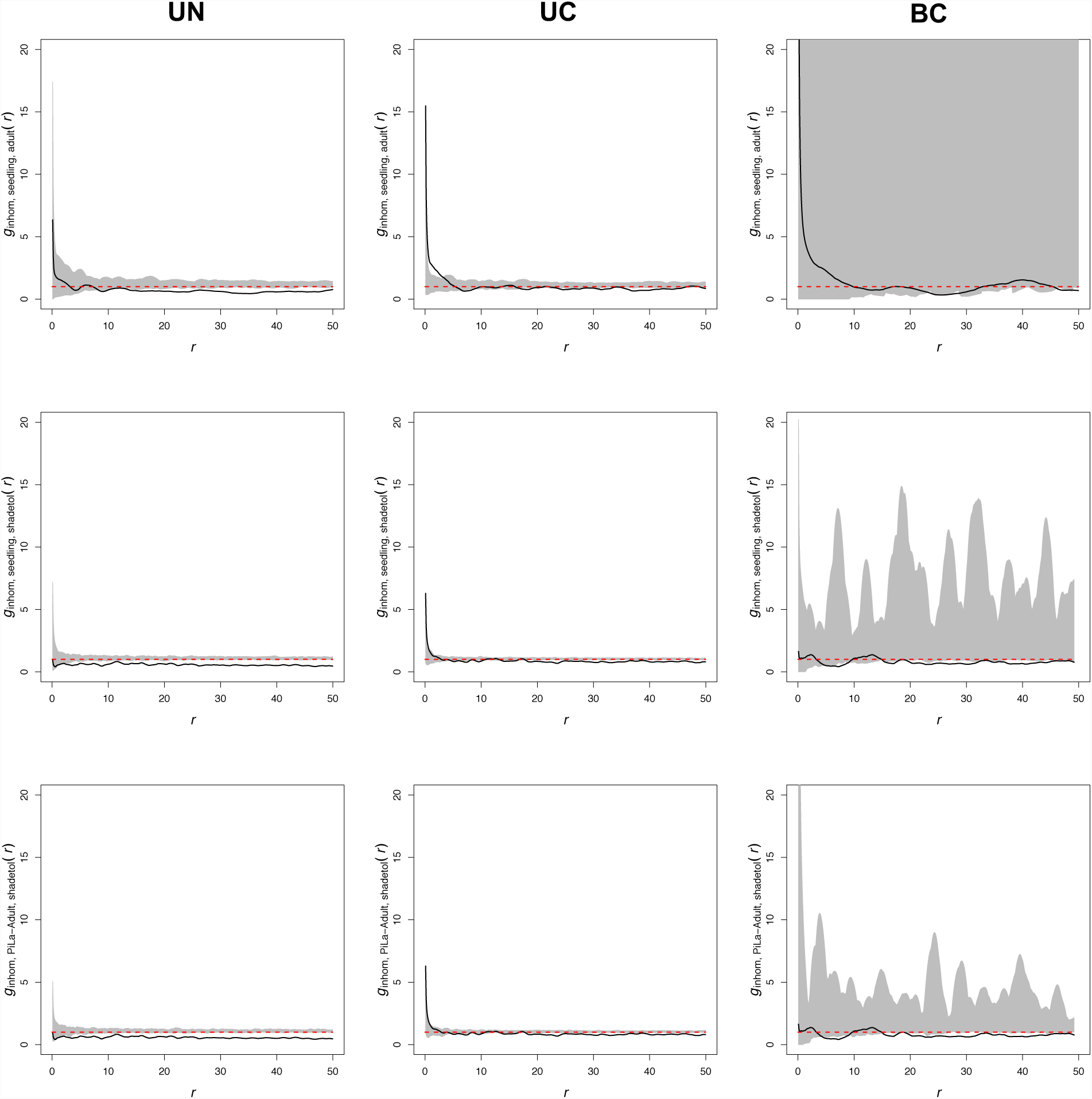
Representative figures of bivariate analysis of spatial structure, *g*_inhom,i,j_(*r*), between: first row: P. lambertiana seedlings (seed) and P. lambertiana adults; second row: P. lambertiana seedlings and shade tolerant adults; third row: P. lambertiana adults to shade tolerant adults. Disturbance intensity increases by column from left to right. These figures show that the two classes compared are generally inhibited spatially by the presence of the other, and that with increasing disturbance there is a diminution of the degree of spatial inhibition between classes. Gray : null confidence envelope; Solid black line : observed *g*_inhom,i,j_(*r*). Red dashed line : null expectation of complete spatial random-ness, *g*_inhom,i,j_(*r*) = 1. Individuals are aggregated if *g*_inhom,i,j_(*r*) > 1, inhibited if *g*_inhom,i,j_(*r*) < 1. The gray shading in the third column of the first row indicates the null confidence envelope extended beyond the limit of the y-axis, where the pattern of the confidence envelope seen in the third column of the second and third rows is caused by sample size varying among distance classes. It should be noted that the observed values for all comparisons generally fall below the y = 1 expectation except for some short distance classes. See Supplemental Figures S4-S6 for all plots.

### Diversity measures

To compare our results with those from the literature we calculated various genetic diversity measures (Table 1) that were most influenced by census size. For instance, census size increased from BC (*n* = 109 individuals) to UN (*n* = 557 individuals) to UC (*n* = 682 individuals) where related diversity measures of *A*_T_, *A*, *A*_*e*_, and *A*_*P*_ followed this trend. Observed heterozygosity was greatest for UN, followed by BC and UC, while expected heterozygosity decreased from UC to BC to UN (Table 1). Thus, no trend was observed between diversity measures and increasing disturbance treatment.

**Table 1.** Genetic diversity measures (standard deviation) by treatment. N: census number of individuals [adults, seedlings]; A_T_ : total number of alleles; A : mean number of alleles per locus; A_e_ : effective number of alleles (harmonic mean across loci); H_o_, H_e_ : respectively the observed and expected heterozygosity for nuclear markers; A_P_ : average number of private alleles. For A, A_e_, H_o_, and H_e_, values indicate averages across loci, where values for each locus were calculated across all three treatment replicates simultaneously. H_o_ and H_e_ used only nuclear markers, whereas other genetic diversity columns considered all loci.

| Treatment | *N* | $A_T$ | $A$ | $A_e$ | $H_o$ | $H_e$ | $A_p$ |
| --- | --- | --- | --- | --- | --- | --- | --- |
| UN | 557 [236,321] | 180 | 25.71 (6.50) | 3.23 (1.58) | 0.87 (0.06) | 0.77 (0.06) | 46 |
| UC | 682 [307,375] | 210 | 30.00 (7.76) | 6.20 (3.07) | 0.57 (0.30) | 0.84 (0.10) | 73 |
| BC | 109 [42,67] | 107 | 15.29 (6.80) | 4.80 (2.46) | 0.82 (0.08) | 0.82 (0.07) | 5 |
| Mean | 449 [195,254] | 165.67 | 23.67 | 4.74 | 0.75 | 0.81 | 41.3 |

Hierarchical *F*-statistics were calculated with nuclear markers to compare the extent of fixation within and across treatments, with individuals nested in replicates, replicates nested in treatments, and treatments nested within TEF. The overall multilocus *F*_ST_ (*F*_*rep,TEF*_) was 0.075, consistent with estimates of many *Pinus* species across various spatial scales (Howe et al. 2003), suggesting that the majority of genetic variation was partitioned more so within plots than between plots. The *F*_*rep,TEF*_ for individual markers varied: rps02 (*F*_*rep,TEF*_ = 0.019), rps12 (*F*_*rep,TEF*_ = 0.037), rps39 (*F*_*rep,TEF*_ = 0.148), rps50 (*F*_*rep,TEF*_ = 0.103). Considering only genotypes across replicates of a given treatment, treatment-level estimates of *F*_rep,tx_ also varied (*F*_rep,UN_ = 0.011, *F*_rep,UC_ = 0.109, *F*_rep,BC_ = 0.035) but showed no pattern with increasing disturbance intensity. Pairwise *F*_rep,tx_ comparisons between treatments were calculated by considering genotypes across two treatments simultaneously and were used to compare the extent of fixation across disturbance intensity. Here, the three comparisons ranged from 0.050 (UC and UN) to 0.055 (UC and BC) to 0.075 (BC and UN) indicative of increasing relative fixation with increasing disparity for the intensity of disturbance for a given comparison.

### Analysis of spatial genetic structure

Analysis of spatial genetic autocorrelation (sensu Smouse & Peakall 1999) was carried out to better understand how treatment affects standing genetic structure (*P. lambertiana* adults x *P. lambertiana* adults), how this standing genetic structure relates to the genetic structure of seedlings (*P. lambertiana* seedlings x *P. lambertiana* seedlings), and the tendency of alike genotypes to be aggregated or inhibited across the treatments as the stands continue to develop (*P. lambertiana* adults x *P. lambertiana* seedlings). In all comparisons, spatial genetic structure in BC treatments did not differ significantly from a random spatial distribution of genotypes (last column Figure 4), perhaps due to the relatively small census sizes (Table 1) in distance-class bins. However, there seems to be an effect of treatment on the spatial patterning of genotypes of adults in the UC and UN stands (first row Figure 4). While UN exhibited small but significant spatial genetic structure for most distance classes up to 200m, UC stands exhibited significant aggregation of adult genotypes at a greater degree than UN up to 150m, where genotypes became spatially inhibited up to the maximum distances in stands 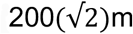; Figure 4). These patterns resulted in spatial distributions of seedling genotypes that were randomly distributed except for very short distance classes in UN, and for UC seedlings, resulted in the general pattern observed for UC adults albeit to a higher degree of both aggregation and inhibition (second row Figure 4). Consequently, alike genotypes between adults and seedlings were aggregated up to 150m in UC, whereas this relationship in UN resulted in negative values of 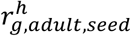 that bordered the confidence envelope but were not significantly different from a random spatial distribution of genotypes (third row of Figure 4). While the genetic structure of adults is due to the interaction of the effect of treatment on pretreatment conditions, the long-term dynamics of these stands will be influenced by seedling ingrowth. These results suggest that UC treatments may, in the long term, increase the relatedness of individuals across short spatial scales less than 150m relative to either BC or UN treatments. This will be particularly exacerbated if gene flow occurs at similarly fine spatial scales (see below).

**Figure 4.**
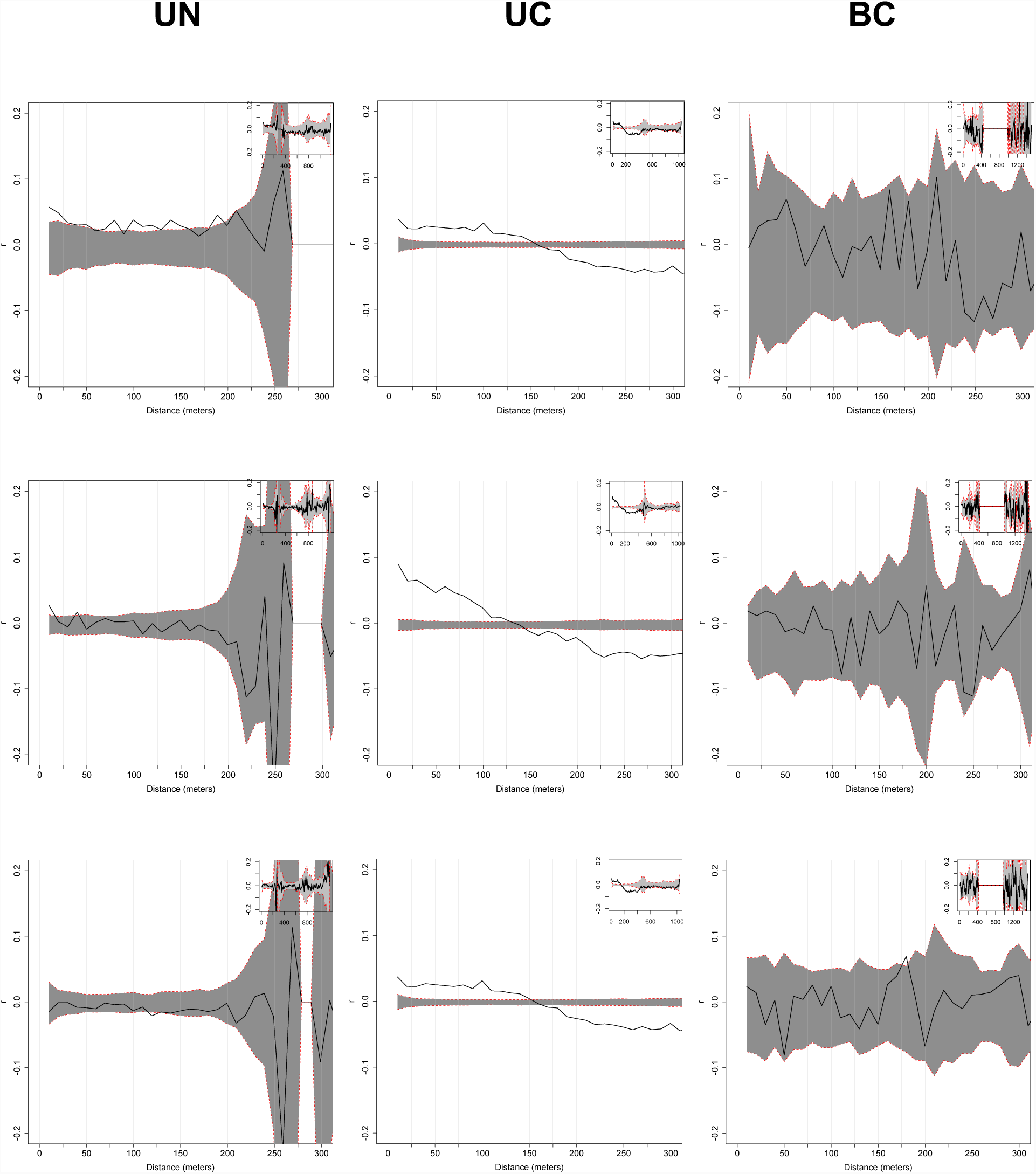
Analysis of spatial genetic structure (sensu Smouse & Peakall 1999) between P. lambertiana adults (first row), P. lambertiana seedlings (second row), and between P. lambertiana adults and seedlings (third row) by treatment (columns) across distance classes within plots (main panel) or across TEF (insets). Values of 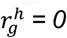 indicate random spatial patterns of genotypes, 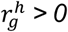 indicate clustering of alike genotypes, and 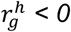 indicate spatial inhibition of alike genotypes.

### Quantifying fine-scale gene flow

#### In-plot vs. out-of-plot dispersal events

To understand how gene flow across plots is influenced by treatment, we quantified the number of in-plot and out of plot dispersal events from pedigrees identified as most probable from our parentage analysis. To account for sample size differences, we calculated the ratio of these values. The number of in-plot and out-of-plot dispersal events between mother and offspring differed by treatment (Figure 5A-B) but not significantly so (*p* > 0.4297). The ratio of these values differed by treatment (Figure 5C), with UC having the greatest proportion of in-plot dispersal events but overall there were no significant differences among treatments (*p* = 0.1926).

**Figure 5.**
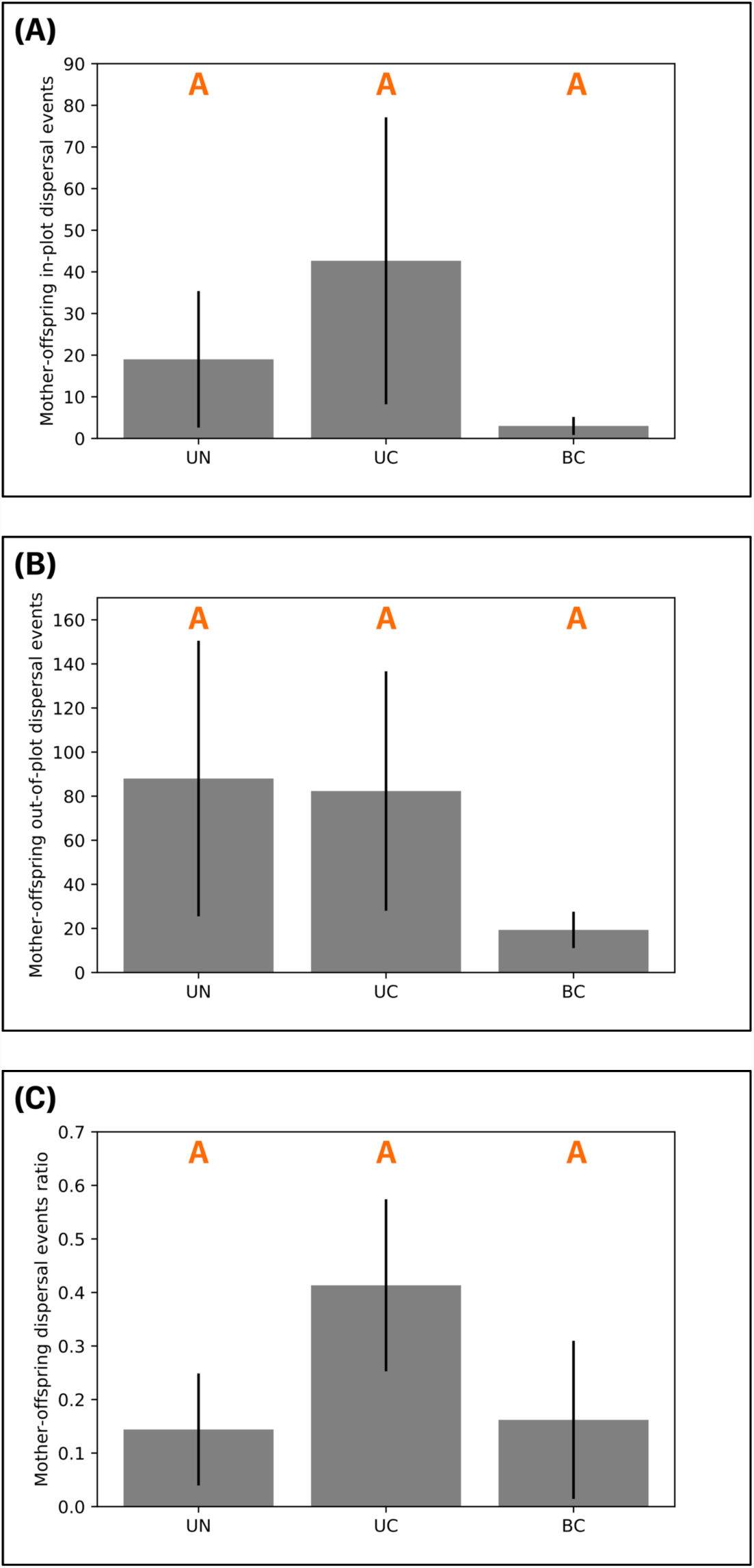
Mother-offspring dispersal events by treatment for (A) dispersal between in-plot individuals, (B) dispersal into plot from an out-of-plot mother, and (C) the ratio of these values. There were no events in which a known mother dispersed seed to another plot, therefore B is utilizing information from parentage analysis that indicated the mother of a given seedling was not sampled. Orange letters within each plot show significant differences between medians, as inferred from separate Kruskal-Wallis tests (see main text of Results). Vertical lines indicate standard deviations.

We next quantified the number of in-plot and out-of-plot dispersal events of pollen from the most probable pedigrees identified from parentage analysis. In these cases, out-of-plot pollen dispersal events were tallied as an in-plot mother receiving pollen from an unsampled or out-of-plot father. The UC treatment exhibited the most in-plot pollen dispersal events, followed by UN and BC (Figure 6A), though not significantly (*p =* 0.5073). UN and UC treatments exhibited similar levels of out-of-plot dispersal events (Figure 6B), which differed (though not significantly, *p* = 0.1376) from BC out-of-plot events. The ratio of in-plot vs. out-of-plot dispersal events increased with increasing disturbance (Figure 6C) but did not differ significantly (*p* = 0.1030).

#### Median dispersal distances by treatment

Considering the most probable parents, we calculated the median seed dispersal distances between offspring and known mothers, and between the median pollen dispersal between known mothers and fathers. Median seed dispersal varied by treatment, being greatest for UN and decreasing with increasing disturbance intensity (Figure 7A). Results indicated significant differences between groups (*p* = 0.0480), with *post hoc* tests indicating significant differences between UN and BC (*H* = 4.34, *p* = 0.0372) but not between UN and UC (*H* = 2.77, *p* = 0.0959) or between UC and BC (*H =* 2.75, *p* = 0.0970; Figure 7A). Median pollen dispersal varied by treatment, being greatest for BC treatments, followed by UN and UC treatments, which did not differ significantly (*p* = 0.1381; Figure 7B).

**Figure 6.**
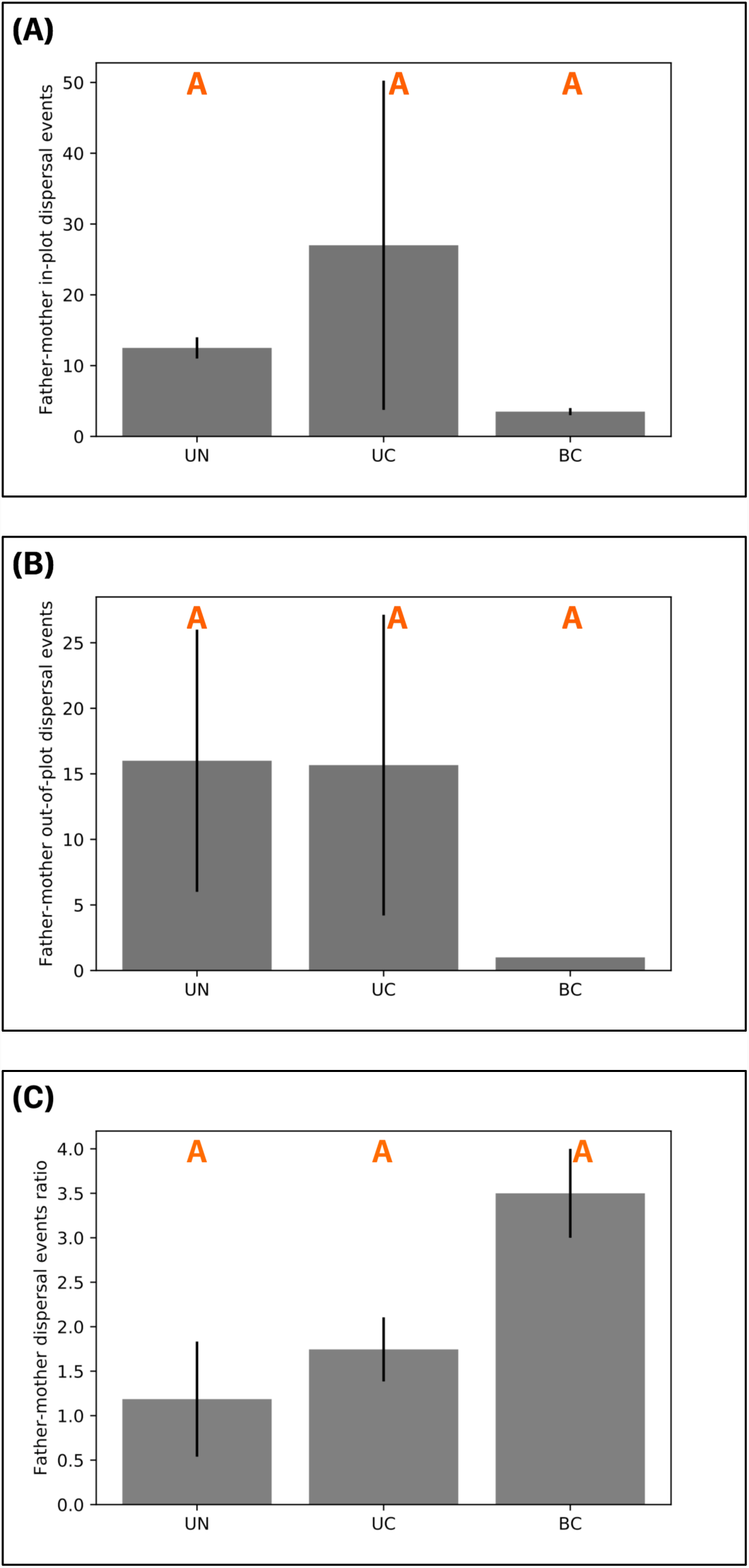
Father-mother dispersal events by treatment for (A) dispersal between in-plot individuals, (B) dispersal into plot from an out-of-plot mother, and (C) the ratio of these values. Plot-level tallies were those of in-plot mothers receiving pollen from either an in-plot father (A) or an out-of-plot (sampled or unsampled) father (B). Orange letters within each plot show significant differences between medians, as inferred from Kruskal-Wallis tests (see main text of Results). Vertical lines indicate standard deviations.

**Figure 7.**
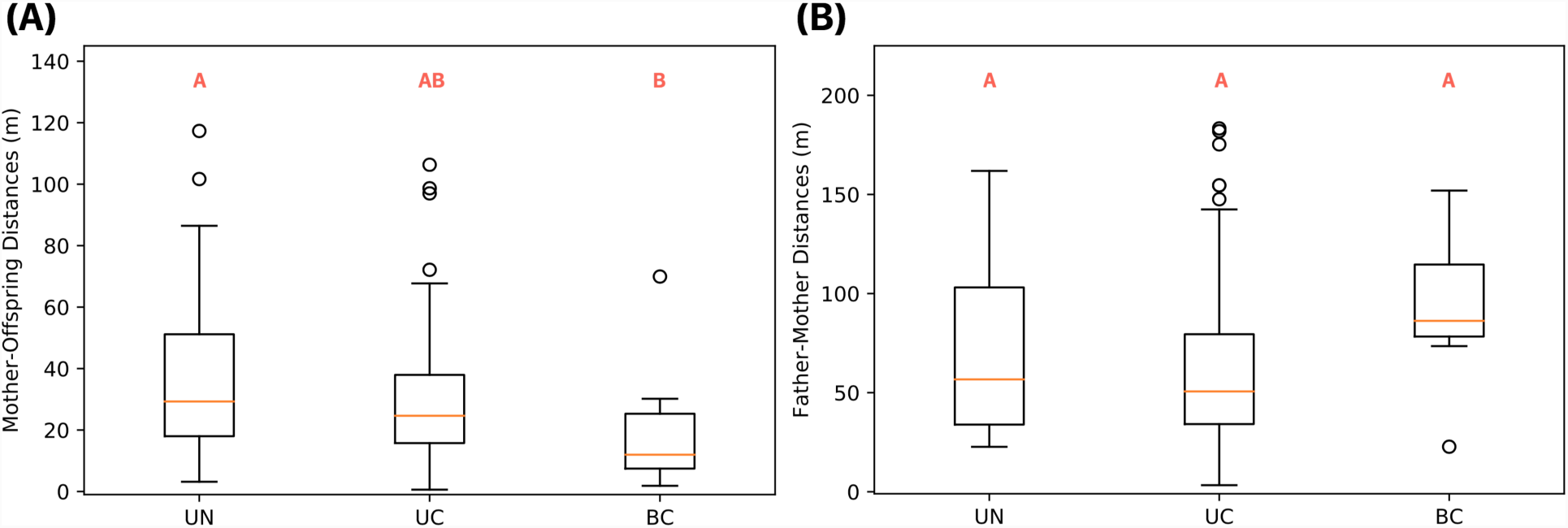
Dispersal distances for seed (A) and pollen (B) calculated from the most probable pedigree from parentage analysis, considering only pedigrees with known mothers (A) or known parents (B). Orange letters within each plot show significant differences between medians, as inferred from Kruskal-Wallis tests for mother-offspring and father-mother dispersal distances (see main text of Results).

These realized distances were roughly in line with mean dispersal distances estimated from dispersal kernel shape parameters in the parentage analysis: mean seed dispersal = 65m (95% credible interval: 57-75); mean pollen dispersal = 170m (95% CI: 150-190; Figure 8).

**Figure 8.**
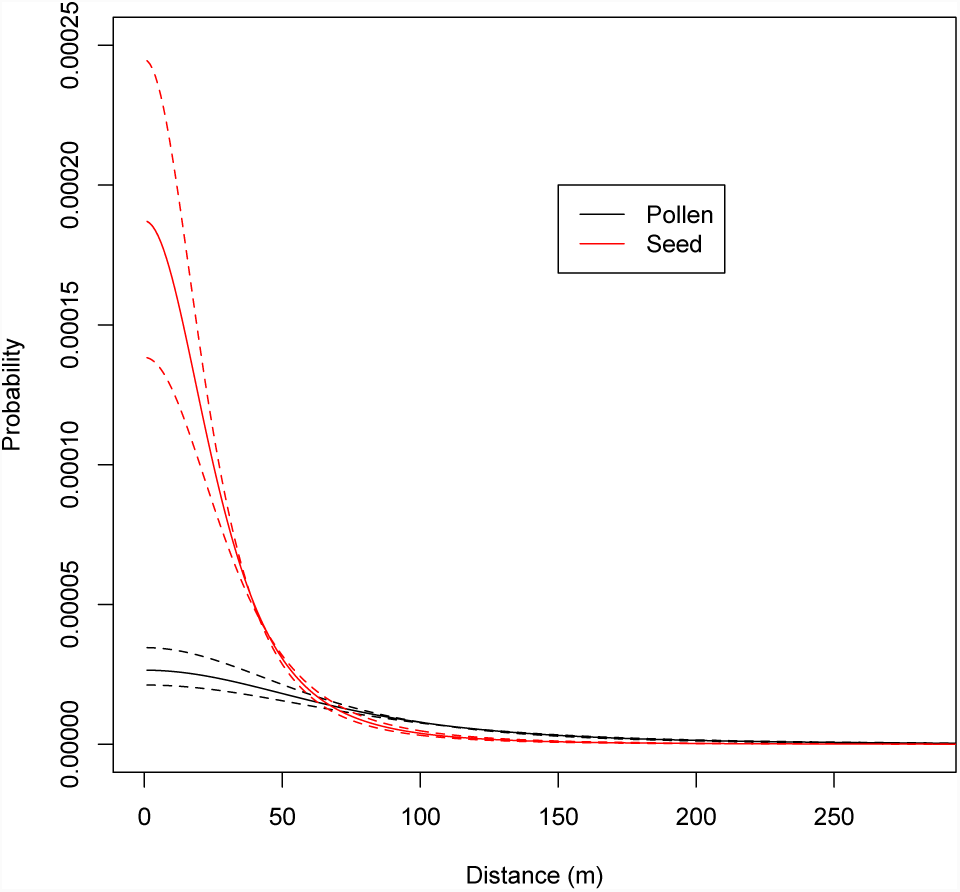
Fitted 2D-t dispersal kernel for seed (red) and pollen (black) using shape parameters inferred from parentage analysis (sensu Moran & Clark 2011). Dashed lines show the 95% credible interval. This figure is truncated at the maximum distance within plots (200√2m) to focus on differences at short distances.

To consider uncertainty in parentage assignment, we calculated weighted average dispersal distances for seed and pollen dispersal. Assignments to mothers of out-of-plot adults were less common than for assignments to in-plot fathers, as can be seen from the blocks (replicates) within treatment of Figure 9. Using fractional parentage, we calculated weighted average distances for each seed and nested these distances within treatments. We first considered mother-offspring and father-mother dispersals from fractional parentage where the identified adults could originate in any treatment. Distances differed significantly by treatment (Figure 10A; *H =* 7.91, *p* = 0.0191) where UN and UC were significantly different (*H* = 8.11, *p* = 0.0044) but not between any other comparison (*H* range = [0.0042,0.6755], *p* > 0.4111). Father-mother distances (Figure 10B) also differed by treatment (*H* = 41.16, *p* = 1.15E-9), with median dispersal distance decreasing from BC to UN to UC, where all pairwise considerations were significant (*H* range = [5.21,27.18], *p* range = [1.85E-07, 0.0224]).

**Figure 9.**
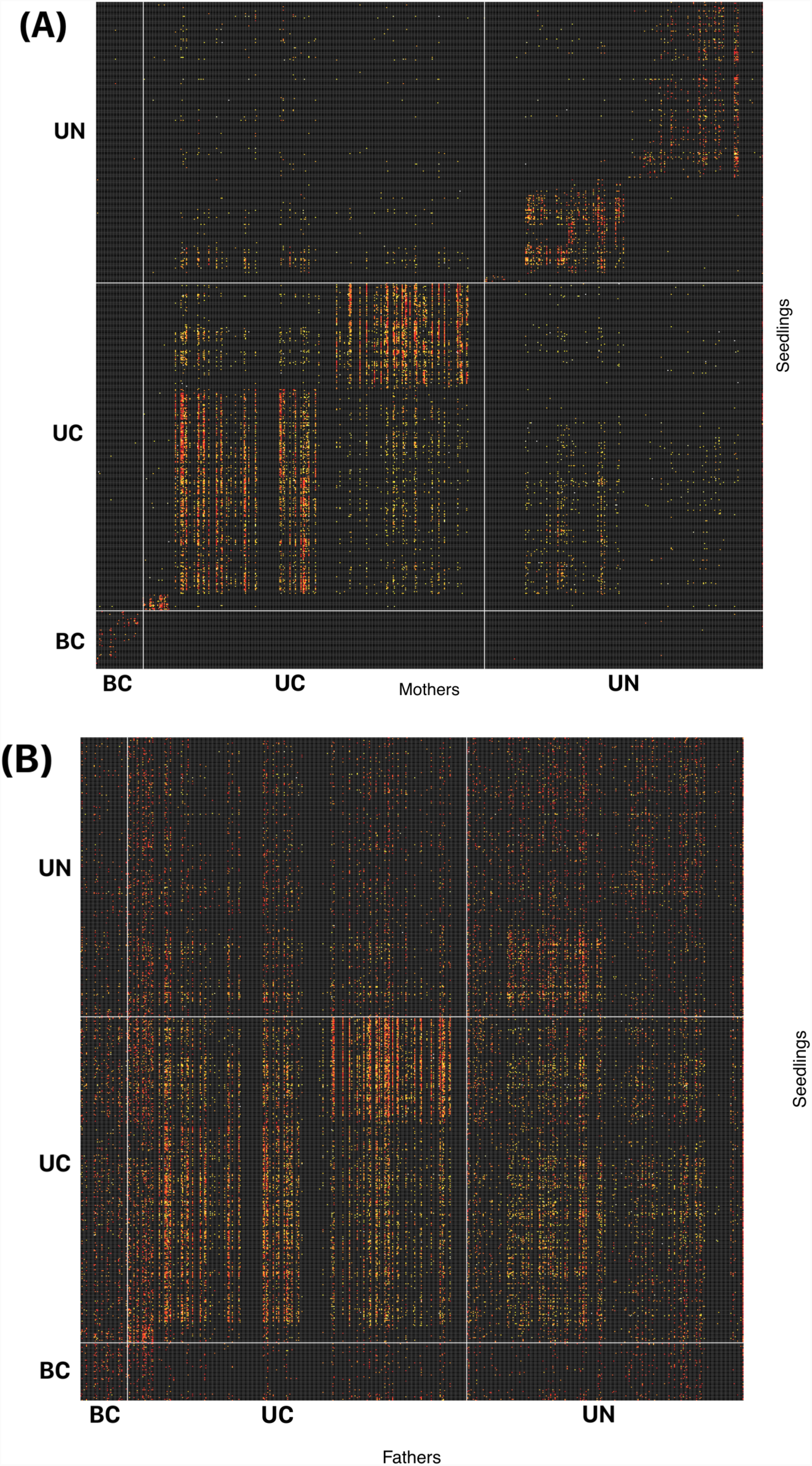
Fractional parentage across parentage analysis cycles for (A) maternal assignment and (B) paternal assignment (see Methods) with adult individuals along x-axes and seedling individuals along y-axes. Each cell represents the fraction of the cycles a particular seedling was assigned to a given adult (black ∼ 0 to red to orange to yellow to white ∼1).

**Figure 10.**
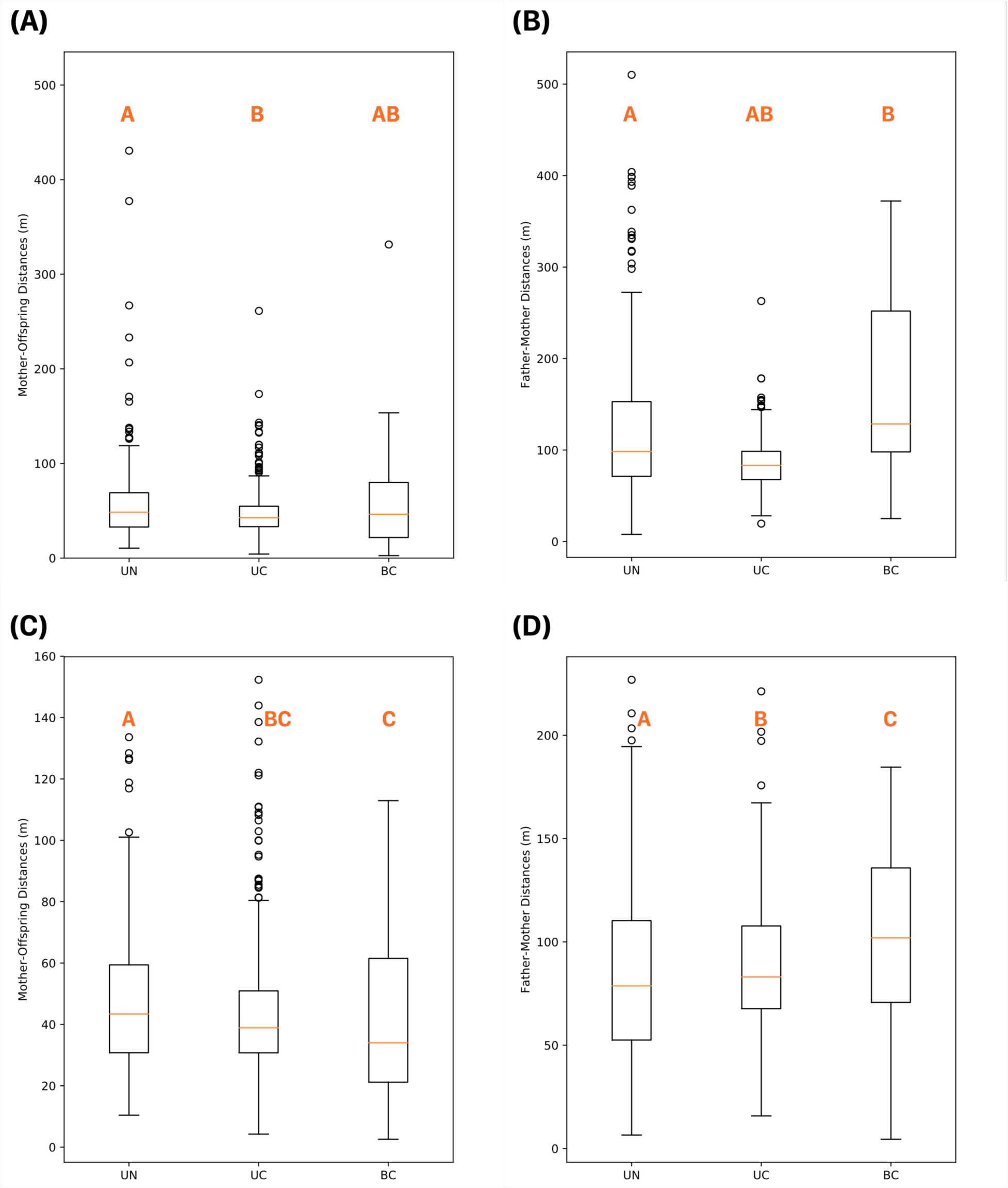
Dispersal distances between mothers and offspring (first column) and between fathers and mothers (second column) using assigned adults from any location (A-B) and for only in-plot individuals (C-D). Orange letters within each plot show significant differences between medians, as inferred from separate Kruskal-Wallis tests (see main text of Results).

Because the proximity of the treatment replicates may interact with dispersal estimates, we considered dispersal distances within plot tallied within treatments using weighted distances. Median values of mother-offspring in-plot distances decreased with increasing disturbance (Figure 10C) and differed by treatment (*H* = 47.10, *p* = 5.91E-11), but only between UN and UC (*H = 4.29*, *p* = 0.0382) and between UN and BC (*H* = 5.83, *p* = 0.0253) and not between UC and BC treatments (*H* = 0.95, *p* = 0.3291). In-plot father-mother distances (Figure 10D) were significantly different across treatments (*H* = 13.89, *p* = 0.0010), with BC having greater distances that either UN (*H* = 5.83, *p* = 0.0157) or UC (*H* = 5.07, *p* = 0.0242), and UC exhibiting greater distances than UN (*H* = 5.00, *p =* 0.0253).

## Discussion

Frequent fires were commonplace in historic forests of the Sierra Nevada, where forests exhibited relatively lower tree densities and a higher proportion of pine species (North *et al*. 2005; Knapp *et al*. 2013). Yet post-settlement fire suppression has led to forest densification that has caused instability in these systems and has increased the chances of uncharacteristic high-severity wildfire. As a result, thinning prescriptions are used to increase the resilience of constituent stands (SNFPA 2004; Agee & Skinner 2005; Schwilk *et al.* 2009; Safford *et al.* 2009). While these prescriptions can mimic the density-reducing effects of fire, and reduce fire severity, it is currently unknown how thinning, in isolation or through its interaction with managed fire, will alter evolutionary dynamics of ecologically important species such as *P. lambertiana* (SNEP 1996). Our results suggest that spatial structure of constituent species is a result of the interaction between treatment and ecology where pines are often clustered with other pines, shade-tolerant trees are often clustered with other shade-tolerant trees, and pine seedlings often are inhibited by both adult pine and shade-tolerant individuals. While genetic diversity statistics are informative of stand-level diversity, they are less informative regarding ongoing evolutionary dynamics as a result of treatment as they do little to predict inbreeding of future generations nor the scale at which mating events are to occur. Used in isolation, diversity indices leave researchers to speculate about ongoing processes and future outcomes, while monitoring of processes that affect fitness provides more meaningful inferences which can be directly used by land managers.

From the analysis of spatial genetic structure (*sensu* Smouse & Peakall 1999), and despite spatial inhibition between adults and seedlings across treatments, our results suggest that unburned thinned stands (UC treatments) result in the increase of fine-scale similarity of adult to seedling genotypes relative to control (UN treatments) or thinned-and-burned stands (BC treatments). Parentage analysis offered additional quantification of fine-scale gene flow and suggested that effective seed and pollen dispersal within plots generally decreased and increased, respectively, with the increasing intensity of disturbance, perhaps due to an increase in microsite suitability for *P. lambertiana* seedlings, or for adults, the availability of potential mates. Our results were measured from individuals remaining or regenerating 13 years post-treatment, very near the historical fire return interval for this area. Thus, ongoing dynamics should be monitored, and will likely change through time, as stands with different treatments continue to develop and respond to subsequent disturbances such as fire.

### The genetic effects of forest management

With some exceptions, studies investigating the genetic consequences of forest management have centered around the impact on genetic diversity indices (see Table 1 in Ratnam *et al*. 2014). This focus is likely due to the fact that highly outcrossing tree species often suffer from elevated inbreeding depression, where survival and reproduction of subsequent generations may be impacted. In such cases, genetic diversity has been used as an index for evolutionary potential, likely attributable to the consequences of the relative contribution of additive genetic variance to phenotypic variance (i.e., narrow-sense heritability) in the breeder’s equation (Lynch & Walsh 1998), but the use of heritability itself as a measure of evolvability comes with important caveats (e.g., see Hansen *et al*. 2011). Further, such diversity indices have been used to assess the relative reduction of alleles due to harvest intensity, where the removal of individuals from stands will likely reduce the diversity of alleles present. Here, management resulting in population bottlenecks is of concern. While these premises are important to investigate, the use of genetic diversity indices as the sole method for inference of management impact are limiting with regard to evolutionary outcomes. If the focus is to be on management impact on evolutionary potential, processes that influence evolutionary fitness should be investigated instead (e.g., mating systems, effective dispersal, fecundity, spatial genetic structure, pollen pool heterogeneity, juvenile survival; Lowe *et al*. 2015). Many traits with fitness consequences in trees are of a polygenic basis (Lind *et al*. 2018), where any given underlying positive-effect locus has minimal influence on the trait. In such cases, fixation (as measured by a handful of putatively neutral markers) at some of the underlying causative loci can be ameliorated by selection for combinations of alleles at other loci. Therefore, while alleles with little to no effect on fitness are informative for demographic processes, these should not be conflated with loci under selection, particularly loci under strong negative selection with important implications for inbreeding depression. Such neutral markers could be better utilized in assessing consequences *within* processes that directly affect fitness. However, in cases where spatial genetic relatedness is increased as a result of management, or individuals become increasingly sparse, wasted reproductive effort (e.g., embryo abortion, or high juvenile mortality) due to increased instances of consanguineous or self-mating events may play an important role in ongoing population dynamics (Woods & Heman 1989; Williams & Savolainen 1996; Sorensen 2001; see also Kärkkäinen *et al*. 1999), particularly when seed rain of heterospecifics exceeds effective reproductive output of historical or ecologically important species (e.g., as is the case for *P. lambertiana* at TEF, Zald *et al*. 2008). Results from tree breeding outcomes also suggest that inbred seeds surviving the embryonic stage will likely have reduced growth and reproductive output at later stages which will also have important consequences to population growth rates and competitive advantages in natural stands (see Rudolph 1981, Sorensen & Miles 1982, Matheson et al. 1995, Durel 1996, Williams & Savolainen 1996, Wu et al. 1998, Petit & Hampe 2001, Savolainen & Pyhäjärvi 2007, Chhatre et al. 2013, Conte et al. 2017, and references therein). For sugar pine in particular, we should expect high inbreeding depression as with most conifers, particularly because of evidence from high diversity and low inbreeding levels found in nearby populations in the Lake Tahoe Basin (Maloney et al. 2011). In addition, genetic diversity will be paramount to the resistance of white pine-blister rust (*Cronartium ribicola*; McDonald et al. 2004).

### Dispersal dynamics of tree species

The analysis of spatial genetic structure and gene flow within and across populations of trees can elucidate ongoing evolutionary dynamics, as this spatial structure is a result of selective and neutral processes acting across temporal and spatial scales (Hardy & Vekemans 1999; Oddou-Muratorio *et al*. 2004; Robledo-Arnuncio *et al*. 2004; Oddou-Muratorio *et al*. 2011). Thus, quantifying dispersal and mating system is an important component in understanding such patterns. There are multiple biological and ecological factors that shape dispersal dynamics and resulting mating systems, such as population density, degree of fragmentation, manner of pollination (e.g., anemophily, entomophily, or zoophily), relative reproductive output, phenotype (such as crown shape or height), interannual climatic variation, as well as stochastic variables such as wind direction and strength (Burczyk *et al*. 1996; Dow & Ashley 1998; Robledo-Arnuncio *et al*. 2004, Burczyk *et al*. 2004; O’Connell *et al*. 2004). Compared with herbaceous and annual plants, trees have more extensive gene flow (Hamrick *et al*. 1992), though such distances are idiosyncratic to a given population, species, and system. For instance, estimates of pollen dispersal for *Pinus sylvestris* varied from between 17-29m based on paternity assignment (Robledo-Arnuncio *et al*. 2004) to 136m (Robledo-Arnuncio & Gil 2005) using the TwoGener method (Smouse *et al*. 2001) where 4.3% of mating events came from pollen dispersed over 30km (Petit & Hampe 2006; Savolainen *et al*. 2007). Seed dispersal distances can also vary idiosyncratically, particularly for winged seeds or those that are also dispersed by animals, such as with *P. lambertiana*.

Spatial genetic structure will be a function of these dispersal consequences as well as their ecological interaction with the environment. While much of the quantification of such structure in trees has been carried out at regional or continental scales, examples exist for investigations at fine spatial scales below a few hundred meters. For instance, Marquardt *et al*. (2007) assessed spatial genetic structure of eastern white pine (*Pinus strobus* L.) as a function of management influence at Menominee Indian Reservation in northeastern Wisconsin. While spatial genetic structure within 100m differed by population, the strongest autocorrelation occurred at the least disturbed site (Marquardt *et al*. 2007). However, while they sampled both adults and natural regeneration they did not distinguish these two groups when inferring spatial genetic structure. Conversely, in Norway spruce (*Picea abies* L. Karst.) populations of northern Italy, Scotti *et al*. (2008) assessed spatial genetic structure of mitochondrial (maternally inherited) and chloroplast (paternally inherited) loci across both adults and saplings. While chloroplast haplotypes were uncorrelated across most distance classes up to 90m for both classes, the maternally inherited mitochondrial markers showed strong affinity below 30m, where this affinity was greater for saplings than for adults. This pattern was seen for *P. lambertiana* individuals at TEF as well, where both adults and seedlings were genetically structured at small distance classes in UC treatments, though seedling genotypes were clustered to a higher degree than adults (Figure 4). To our knowledge however, few instances in the literature compare both spatial structure of trees with spatial genetic structure of tree genotypes. At TEF, seedlings were clustered at fine spatial scales across all treatments likely due to microsite suitability (as most cached seeds will likely persist only in suitable sites), but were only clustered genetically in UC treatments. As such, without genotypic data, investigators may be lead to spurious conclusions where it may be assumed that clustering of individuals also indicates clustering of genotypes. Further, ingrowth of *P. lambertiana* in UC treatments will likely be more related to nearby individuals, which may cause inbreeding and embryo abortion to a greater degree in subsequent generations than in other stands at TEF.

### Management implications

Our results suggest that management is affecting dispersal through the availability of suitable microsites for seedling establishment, as well as through the availability of mates. As disturbance intensity increased at TEF, mean effective seed dispersal generally decreased while effective pollen dispersal generally increased (Figure 7A-B), likely due to the proximity of suitable (e.g., unshaded) microsites and the availability of potential mates, respectfully. Using the inferred dispersal kernels (Figure 8), the vast majority of dispersal occurs across small distance classes, with the estimated probability of dispersal of pollen below 150m accounting for more than 90.2% of pollen dispersal events, while dispersal of seed below 50m (150m) accounts for 87.3% (99.2%) of dispersal events across TEF. Such a dispersal tendency will drive spatial genetic structure and will interact with environment (including management) to ultimately determine the patterns we observe across the landscape. Because UC treatments generally resulted in an increased spatial affinity of alike genotypes between adults and seedlings (Figure 4), short-term dynamics (decadal scales) may be dominated by mating events between related individuals. However, long-term dynamics will likely affect this structure as well. The strong levels of spatial genetic structure observed in seedlings have been shown to decrease in adult stages because of self-thinning processes in other tree species (Hamrick *et al*. 1993; Epperson & Alvarez-Buylla 1997; Chung *et al*. 2003; Oddou-Muratorio *et al*. 2004), and may well occur at TEF as well. Even so, such consequences are dependent upon initial structure that may vary to differing degrees in undisturbed stands, or across the landscape. Long-term dynamics should be monitored as these stands continue to develop and respond to contemporaneous ecological pressures.

## Conclusion

Understanding how thinning and fire prescriptions intended to decrease fire severity and restore ecosystem resilience influence evolutionary dynamics of historically dominant and ecologically important pine species is of paramount significance. We found that treatment of fire-suppressed populations of *P. lambertiana* differentially affects fine-scale spatial and genetic structure, and that seed and pollen dispersal increase and decrease, respectively, with disturbance intensity. Such dynamics are likely to remain unequilibrated in the short term, and therefore management would benefit from further monitoring of evolutionary dynamics that affect fitness in these forests (e.g., reproductive output, survival of seedlings). Further monitoring across broader spatial scales would also inform how these management prescriptions affect dynamics across a greater extent of environmental heterogeneity and how these evolutionary dynamics vary by locality. Such information will allow management to prescribe treatments in a regionally- and site-specific manner.

## Acknowledgments

The authors wish to thank Britta Austin, Casey Harless, Erin Hobson, Nate Stearrett, and Alexandrea Stylianou for help with field and lab work (and for being great people), VCU’s Center for High Performance Computing, VCU for start-up funds awarded to AJE, and helpful comments from two anonymous reviewers.

